# Report of mutation biases mirroring selection in *Arabidopsis thaliana* unlikely to be entirely due to variant calling errors

**DOI:** 10.1101/2022.08.21.504682

**Authors:** J. Grey Monroe, Kevin D. Murray, Wenfei Xian, Pablo Carbonell-Bejerano, Charles B. Fenster, Detlef Weigel

## Abstract

It has recently been proposed that the uneven distribution of epigenomic features might facilitate reduced mutation rate in constrained regions of the *Arabidopsis thaliana* genome, even though previous work had shown that it would be difficult for reduced mutation rates to evolve on a gene-by-gene basis. A solution to Lynch’s equations for the barrier imposed by genetic drift on the evolution of targeted hypomutation can, however, come from epigenomic features that are enriched in certain portions of the genome, for example, coding regions of essential genes, and which simultaneously affect mutation rate. Such theoretical considerations draw on what is known about DNA repair guided by epigenomic features. A recent publication challenged these conclusions, because several mutation data sets that support a lower mutation rate in constrained regions suffered from variant calling errors. Here we show that neither homopolymer errors nor elevated mutation rates at transposable elements are likely to entirely explain reported mutation rate biases. Observed mutation biases are also supported by a meta-analysis of several independent germline mutation data sets, with complementary experimental data providing a mechanistic basis for reduced mutation rate in genes and specifically in essential genes. Finally, models derived from the drift-barrier hypothesis demonstrate that mechanisms linking DNA repair to chromatin marks and other epigenomic features can evolve in response to second-order selection on emergent mutation biases.

## INTRODUCTION

A consortium that included a subset of the authors of the current study (J.G.M., P.C.-B., C.B.F., D.W.) recently reported a link between mutation rates and local epigenomic features enriched in gene bodies and evolutionarily constrained genes [1], with variation in mutation rate probability paralleling patterns of neutral and functional polymorphism in natural populations of *Arabidopsis thaliana.* The epigenomic feature most strongly associated with lower mutation rate was H3K4me1, whose deposition in plants is coupled to transcription and which is enriched in gene bodies and functionally constrained genes [2]. We demonstrated with simulations how epigenomic features that are characteristic for broad genomic regions, such as constitutively expressed genes, and which simultaneously affect mutation rate, could yield a solution to Lynch’s equations for the barrier imposed by genetic drift on the evolution of targeted hypomutation [3–5]. Using data from mutation accumulation lines we derived a genome-wide model of mutation probability, which indicated that epigenome-driven bias can reduce the occurrence of deleterious mutations in *A. thaliana* [1]. This conclusion aligns with an increasing body of mechanistic knowledge of localized DNA repair and its effects on local mutation rate [6–12].

A recent publication [13] pointed out that several of the data sets in [1], especially those that were used to empirically validate mutation probability predictions, did not sufficiently account for variant calling and sequencing errors, in particular bleed-over errors at sites adjacent to homopolymers. A related concern is whether the observations in [1] were confounded by other features, including increased mutation rates in transposable elements (TEs) as a result of elevated cytosine deamination [14]. Here, we show that the findings from [1] are robust to several sources of sequencing errors, that they align with mutation bias observed in a very large rice fast-neutron mutation data set, and that they are supported by a new meta-analysis of a range of *A. thaliana* germline mutation data sets. We discuss the findings in the context of rapidly accumulating evidence for epigenome-guided DNA repair being particularly effective at genes and especially at essential genes.

## RESULTS

In the original study [1] critiqued by Liu and Zhang [13], we had made use of both germline mutation calls and putative somatic mutation calls. For the germline mutations, we only considered homozygous mutations, as in previous mutation accumulation studies [15–21], since *A. thaliana* is selfing and heterozygous mutations will rapidly become either fixed or lost. Homozygous germline mutations can generally be called with high confidence using relatively simple filters. On the other hand, somatic mutations are highly unlikely to be homozygous and in addition most will be restricted to a subset of tissue analyzed. Confident identification of somatic mutations from sequencing data of entire plant rosettes is therefore much more difficult, and even more so when samples originate from pooled individuals. While the mutation probability model from [1] relied to a large extent on homozygous germline mutations, empirical confirmation of predicted patterns was overwhelmingly done with putative somatic mutations. Here we first address the potential influence of homopolymer errors, which is particularly relevant for somatic mutation calls [13], and of increased mutation rate in TEs due to cytosine deamination, which is relevant for both germline and somatic mutations.

### Homopolymer (polynucleotide) errors cannot explain observed mutation biases

Homopolymeric sequences, defined as consecutive runs of identical nucleotides, pose specific challenges for the discovery of spontaneous mutation. On the one hand, homopolymers are particularly likely to produce sequencing errors at immediately neighboring nucleotides through homopolymer bleed-through [22]. On the other hand, homopolymers have much higher true mutation rates in many species [23–28]. They can expand and contract due to uncorrected replication polymerase slippage errors, which are otherwise repaired by mismatch repair, and their distribution across genomes could thus reflect differential efficacy of repair pathways. As such, the heterogeneity in homopolymer density described in [13] could itself be a product of the non-uniformity in DNA repair activity across the genome.

To explicitly address the potential confounding effects of homopolymer errors, we revisited the non-homozygous variant calls that had been first reported in [16] and then reanalyzed in [1] to identify putative somatic mutations. These somatic mutation calls were used together with germline calls, which were essentially the same as in [16], to build a genome-wide model of mutation probability as a function of genomic and epigenomic features [1]. Specifically, here we remapped and called variants against a new, more complete reference genome assembly, to reduce issues arising from mismapping [29]. We then removed calls potentially resulting from homopolymer sequencing bleed-overs and used more stringent filtering criteria, including read support from both strands, alternative calls supported by reads in only one sample, mapping quality ≥ 60, unclustered distance >10 bp to neighboring variants, and location within 3 kb of gene bodies. This more stringent approach resulted in 2,661 instead of 4,317 somatic single-nucleotide substitutions. The mutational patterns observed remained consistent with the original findings from [1], with mutation rates being lower in gene bodies (Fig. 1a). We also note that essential genes contain a greater density of homopolymers in coding regions compared to other classes of genes, so erroneous calls near homopolymers should therefore increase, not decrease apparent mutation rate in coding regions of essential genes.

**Figure 1.**
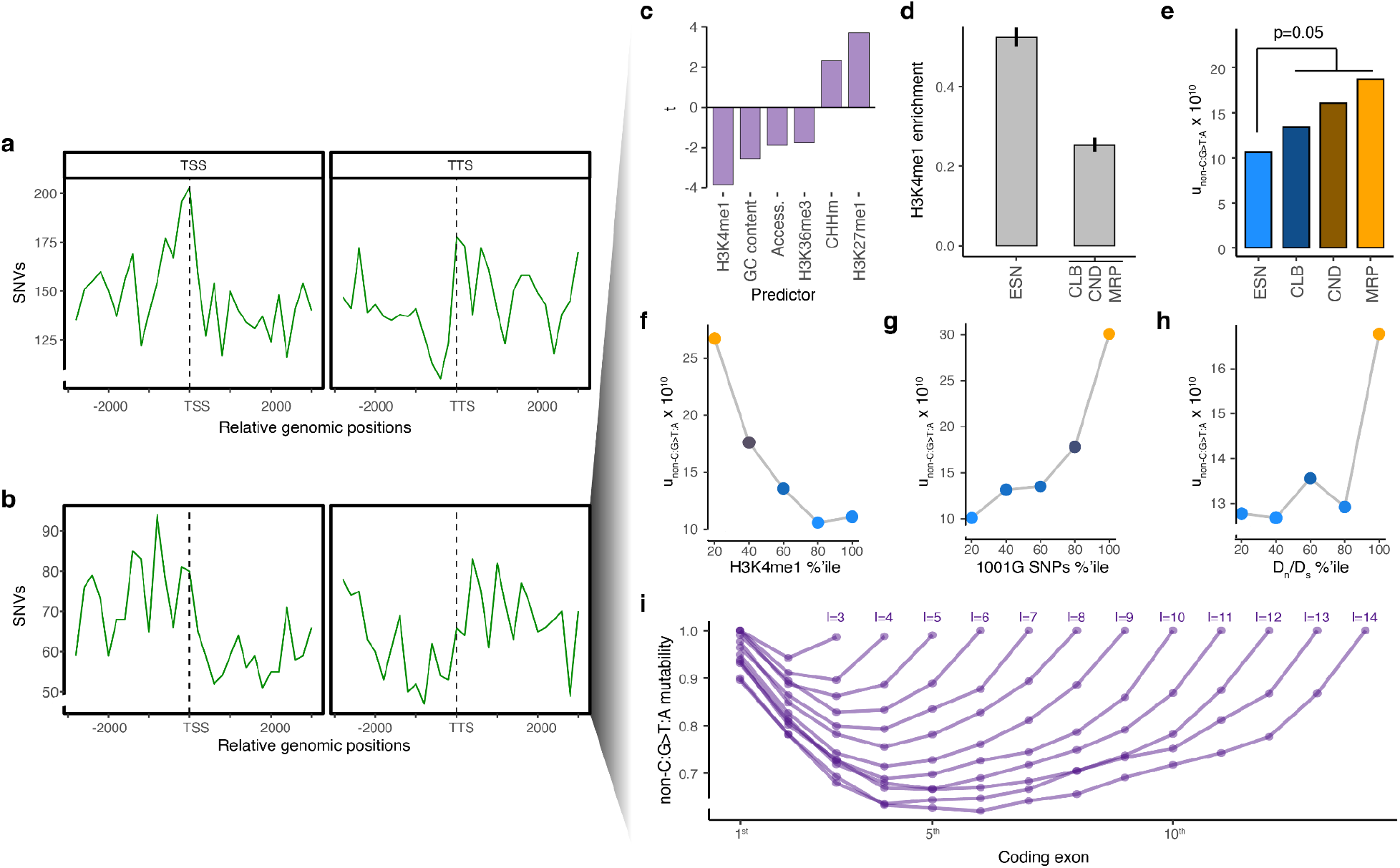
Primary conclusions of [1] are robust to potential variant calling artifacts. **(a)** Genomic sequencing reads of mutation accumulation lines that constitute the main dataset of [1] were mapped to a more complete reference genome, and potential somatic variants were filtered more stringently, as described in the main text. Mutations (shown in 300 bp windows) are reduced in gene bodies, as observed in the original report [1]. TSS, transcription start site; TTS, transcription termination site. **(b)** Distribution of homozygous germline mutations from [16] and 400 new mutation accumulation lines from [1], excluding all C:G>T:A mutations. Further analysis of this reduced germline data set confirms: **(c)** Mutation rates are predicted by epigenomic features, especially H3K4me1; **(d)** H3K4me1 is more highly enriched in gene bodies of essential genes (ESN) than of genes with cellular and biochemical mutant phenotypes (CLB) or environmentally conditional mutant phenotypes (CND) or morphological mutant phenotypes (MRP); **(e)** Mutation rates are lower in gene bodies of essential genes; **(f)** Mutation rates are lower in genomic regions marked by H3K4me1; **(g)** Mutation rates are correlated with rates of genetic diversity in natural populations from the 1001 Genomes Project [30]; **(h)**, Mutation rates are associated with D_n_/D_s_ (ratio of non-synonymous to synonymous substitutions between *A. thaliana* and *A. lyrata)* as an indicator of purifying selection; **(i)** Modeled mutation rates are lower in internal exons. Results were similar for all germline single nucleotide variants (SNVs) or when excluding coding regions to take cryptic selection into account.

In [1], two additional datasets of somatic mutation calls were used for empirical validation of lower mutation rates in gene bodies and in essential genes, one from a new collection of mutation accumulation lines [1], and one that used available sequencing data from a deeply sequenced sets of 64 leaves and that had been used before to identify very-high-confidence somatic mutations [31]. A pipeline used only for the latter dataset resulted in a very large number of raw variants. We since found that reads had inadvertently been mapped twice to the genome. This led to singletons being called as potential non-singleton variants, thus blurring the line between sequencing errors and low-frequency somatic variants (cf. Extended Data Fig. 4c in [1]). We thank the colleagues who pointed out this mistake; corrected analyses that exclude the erroneous calls can be found in the Supplement.

In [1], we had explored different filtering criteria for somatic mutations. Because we had found qualitatively similar patterns in the combined analyses of germline and somatic mutations, we maximized the number of mutations in several summary figures by using rather loose filtering criteria. Specifically, we initially used 2,257 germline and 6,317 stringently filtered somatic mutations (single-nucleotide substitutions and small indels) from the original study of 107 mutation accumulation lines [16] to build a model of epigenome-driven mutation rate bias. For the empirical confirmation of the observed reduced mutation rates in gene bodies and essential genes, we had added in [1] 8,891 germline mutations from a natural mutation accumulation line experiment [32], 3,306 germline and 355,827 loosely filtered somatic mutations from 400 additional mutation accumulation lines first reported in [1], and 773,141 somatic mutations from 64 leaves (which included a large number of singletons that were erroneously called as variants, as discussed in the preceding paragraph) [31]. For context, DNA is damaged 10,000 to 100,000 times per day in each cell, at least in humans, with cases of unrepaired damage going on to become true *de novo* mutations [33–35]. A recent analysis of human somatic mutations found 484,678 single base pair mutations in 389 healthy samples sequenced at 27x coverage [36], and another study reported up to ~7 mutations/Mb in healthy human cells [37]. Based on empirical measurements and by extrapolating from relationships between life history and somatic mutation rates, somatic mutation rates in *A. thaliana* could be many orders of magnitude higher than in humans [38,39]. Liu and Zhang [13] conclude that the data in [1] imply mutation rates that are around 2 x 10^-4^ bp^-1^ per generation, but this confounds germline and somatic mutations. Because we did not sequence individual somatic cells, it is difficult to estimate their mutation rate but a back-of-the-envelope calculation of ~12,000 sequenced genomes in the 400 additional mutation accumulation lines would yield a rough estimate of a lower bound of 3 x 10 ‘ bp^-1^ per somatic cell. A similar calculation would yield a rough estimate of 10^-5^ bp^-1^ per somatic cell for the 64-leaves data set.

### Additional potential confounders cannot explain observed mutation biases

A confounder when studying mutation bias comes from the observation that TEs, which are more frequent in intergenic regions, are often highly methylated at cytosines, which can be spontaneously deaminated to thymine [14]. A scenario that we did not explicitly test in [1] was whether spontaneous deamination of methylated cytosines in intergenic TEs might be primarily responsible for an increase in mutation rate outside genes. We therefore removed all C:G>T:A mutations from the single-nucleotide germline mutations in [1], which left 1,680 out of 3,550 variants. Even with the reduced mutation set, we still find fewer mutations in gene bodies and in essential genes (Fig. 1b, d, e). We continue to see complex patterns of mutational variation, including an increase in predicted mutability in peripheral exons (Fig. 1i), which is mirrored by patterns of sequence evolution (Fig. 1g, h) (cf. Extended Data Fig. 7 from [1]). We also continue to observe a robust association between lower mutation probability and H3K4me1 (Fig. 1c, f), which is enriched in gene bodies and essential genes. This is consistent with targeting of plant DNA repair proteins via Tudor domains to H3K4me1 [40], as discussed in more detail below.

TEs are also enriched in pericentromeric regions, which are particularly likely to suffer from erroneous variant calling because of their highly repetitive nature. However, concerns pertaining to this issue would not apply to the analyses in [1], since only variant calls near genes (gene bodies plus 3 kb upstream and downstream) were considered, which led to exclusion of gene-poor peri-centromeric regions.

A more significant concern in general is that somatic mutation calls, especially when loosely filtered, as in the validation data sets of [1], are much less reliable than homozygous germline mutations. The main conclusion of [1] did, however, not solely rely on somatic mutation calls. Instead, it was based on combining insights from germline mutations, population genetic variation, mechanisms of epigenome-driven mutation repair, and modeling the compatibility of the drift-barrier hypothesis with mutation rate evolution. In addition, the genome-wide mutation probability models that we developed were very similar when we used only germline mutations, or excluded genic sequences, as described in [1]. The validation sets used to test the predictive power of the mutation probability patterns with empirically observed mutations (Fig. 2b, 3c, and 4c in the main text of [1]) were, however, overwhelmingly from putative somatic mutation calls. The uncertainty of somatic mutation calls should have been incorporated into the statistical tests in [1], and the substantial weight of putative somatic mutations in the empirical confirmation should have been made clearer in [1].

**Figure 2.**
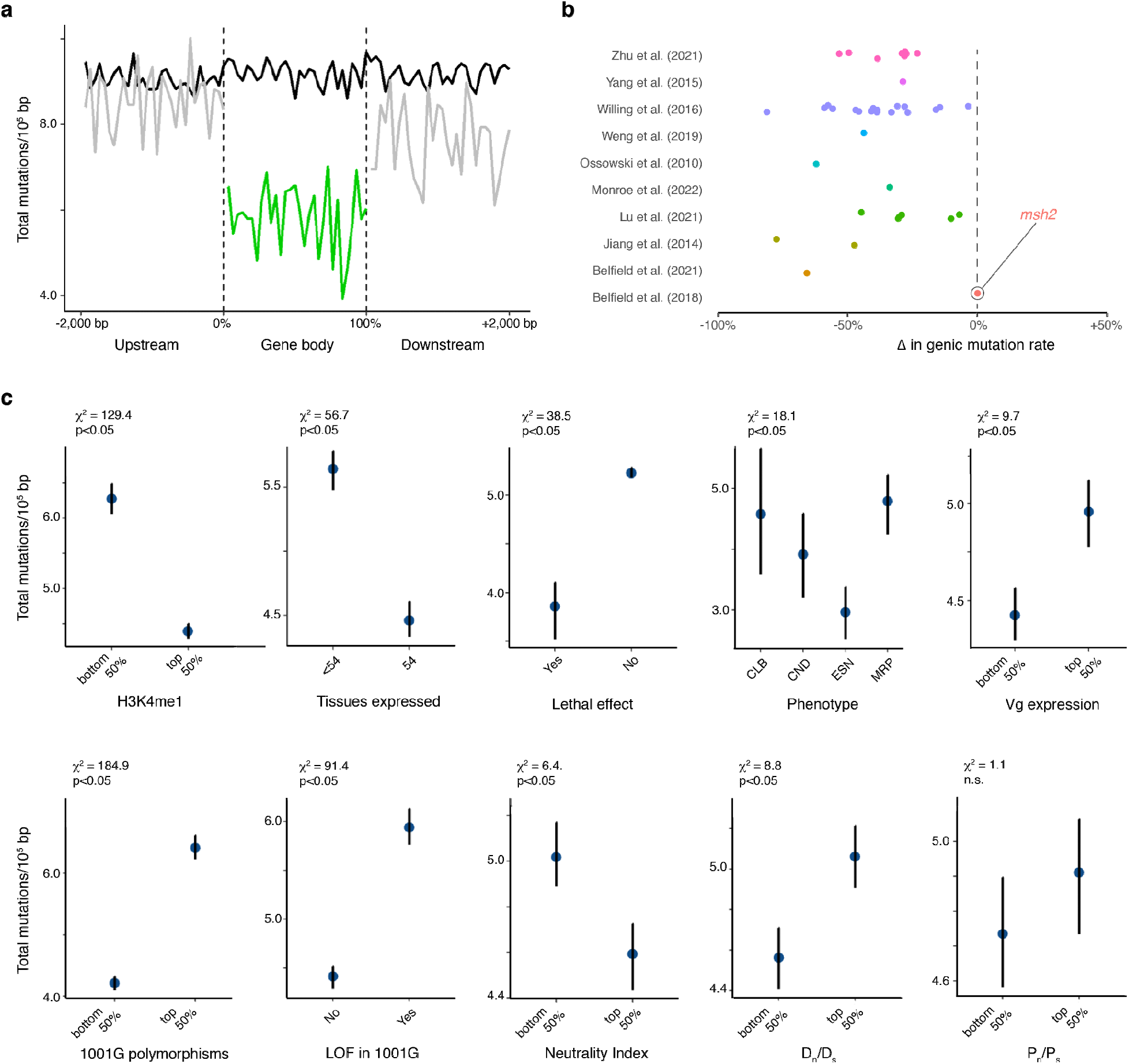
Joint analyses of germline mutations in several published *A. thaliana* mutation accumulation experiments. **(a)** Mutation rates outside (gray lines) and inside gene bodies (green line). The black line indicates random windows in the genome with lengths that reflect the distribution of gene lengths. **(b)** Reduction in genic single-nucleotide mutation rates compared against genomic background, as reported in germline mutations from multiple *A. thaliana* data sets (Table 1). Y-axis values are randomly “jittered” to visually distinguish data from different experiments in the same study. For the study by Monroe and colleagues [1], only new germline mutations from 400 mutation accumulation lines are shown; the other germline mutations in that paper were already described by Weng and colleagues [16], shown separately here. The mutation rate reduction in genic regions is eliminated in *msh2* DNA repair mutants [10]. **(c)** Mutation rates in genes classified by functional category, rates of sequence evolution, patterns of expression, and estimates of selection. Error bars reflect 95% confidence intervals from bootstrapping. Data sources are [1,15–17,19–21,41,42]; see Table 1 for details.

**Table 1.**
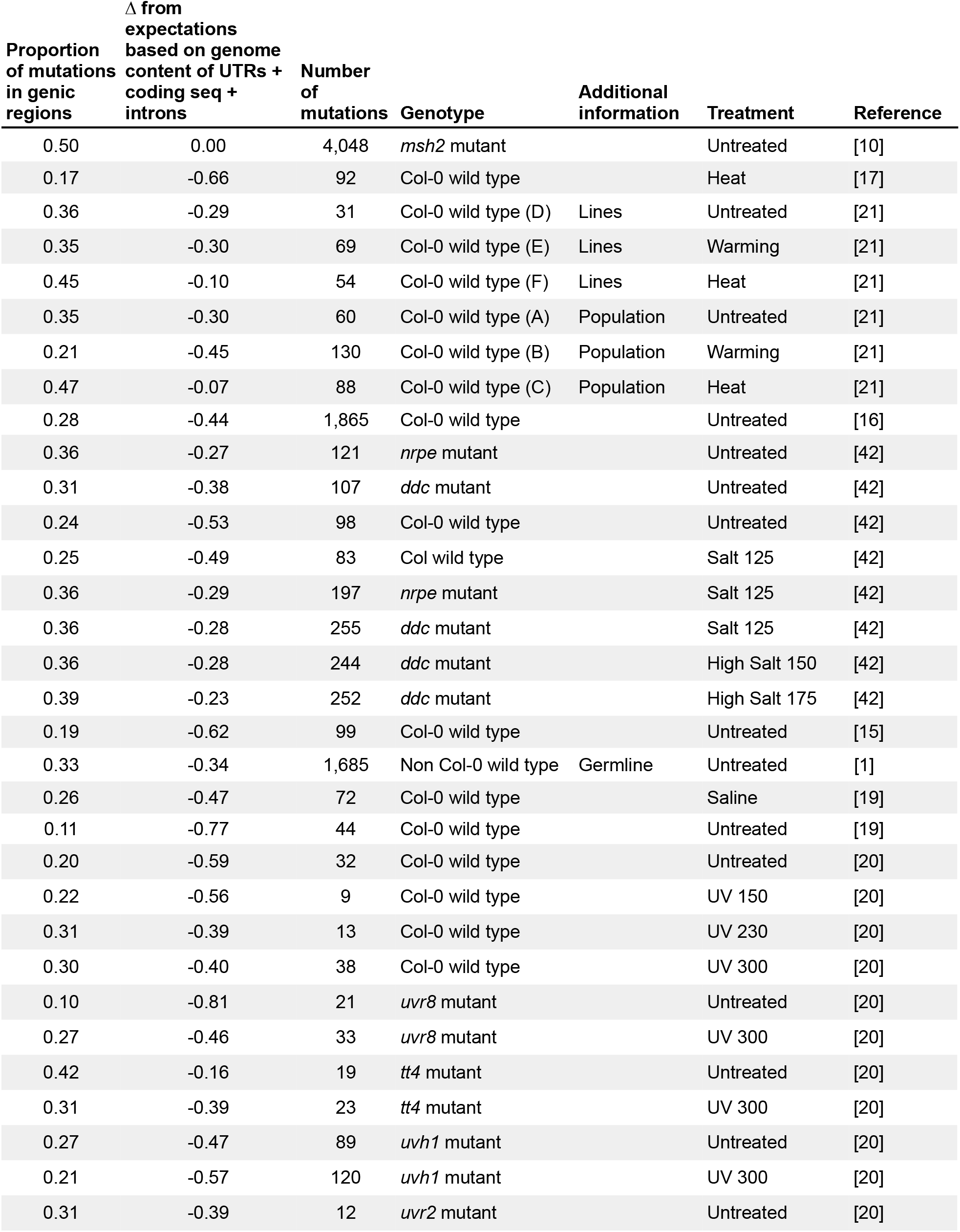
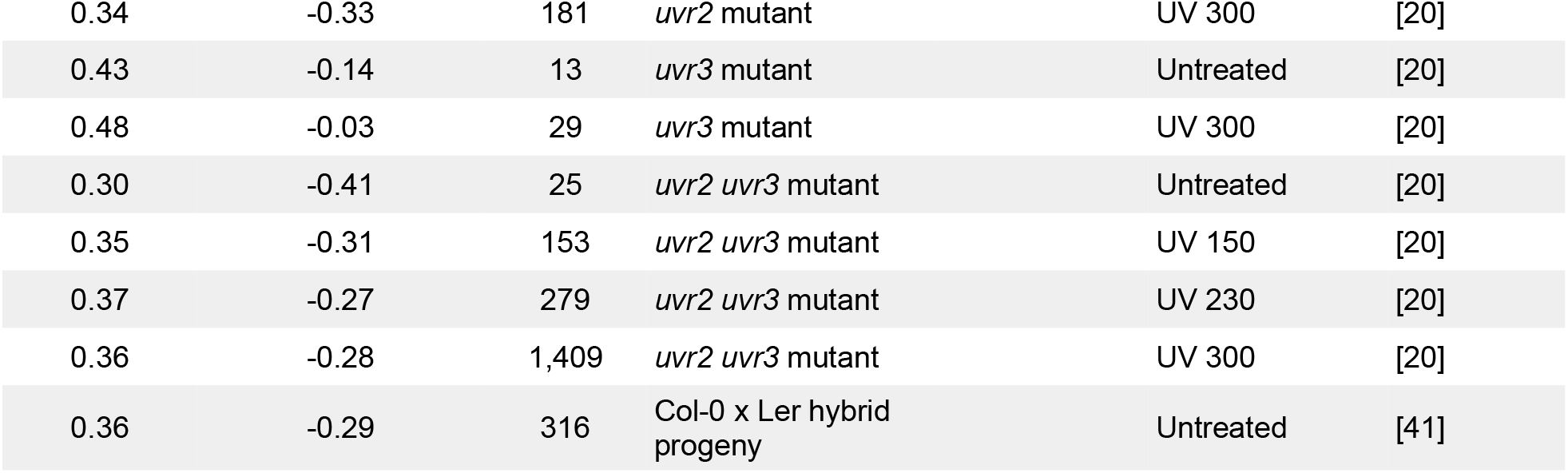
Relative genic mutation rate (only single-base-pair substitutions) in multiple data sets.

### Additional datasets confirm that germline mutation rates are lower in constrained genes

Given the inherent uncertainty of putative somatic mutation calls, we conducted additional analyses that excluded all somatic mutations from the data sets used in [1]. To this end, we combined germline mutations from [1] with results from mutation accumulation experiments in *A. thaliana* generated by several other labs [15–17,19–21,41,42] (Table 1). We did not recall variants, but directly used the mutation calls, the vast majority of which were homozygous calls, as reported in those papers. As had already been described in several of the individual reports, our new meta-analysis revealed a nearly-universal reduction in single-nucleotide mutation rates in gene bodies (Fig. 2a, b, Table 1). The notable exception is in plants lacking the mismatch repair protein MSH2 [10] (Fig. 2b), consistent with DNA repair being targeted to gene bodies, especially of active genes, as demonstrated extensively in humans [6,8–10]. The finding of elevated mutation in gene bodies of *msh2* mutants, in which over 8,000 mutations were mapped, suggests that selection is not the main driver of reduced genic mutation rates detected in the other data sets. The association of H3K4me1 with reduced mutation rate described in [1] is confirmed in this meta-analysis as well: gene bodies, constitutively expressed genes, essential genes, and genes exhibiting evidence of purifying selection in natural populations all have lower germline mutation rates (Fig. 2c).

The patterns around genes from [1] are also confirmed in a reanalysis of a recently published set of 43,483 very high-quality germline variants obtained after fast-neutron mutagenesis in rice and validated at 99% accuracy for single-nucleotide variants [43] (Fig. 3). Mutation rates in this dataset are systematically lower in gene bodies and regions marked by H3K4me1, including active genes and those subject to stronger purifying selection in populations. These findings are not reasonably explained by selection affecting mutation accumulation – mutation rates are lower in genes under purifying selection even when one considers only synonymous mutations. As in *A. thaliana*, removal of C:G>T:A single-nucleotide mutations does not greatly change the observed patterns. More details can be found in [44].

**Figure 3.**
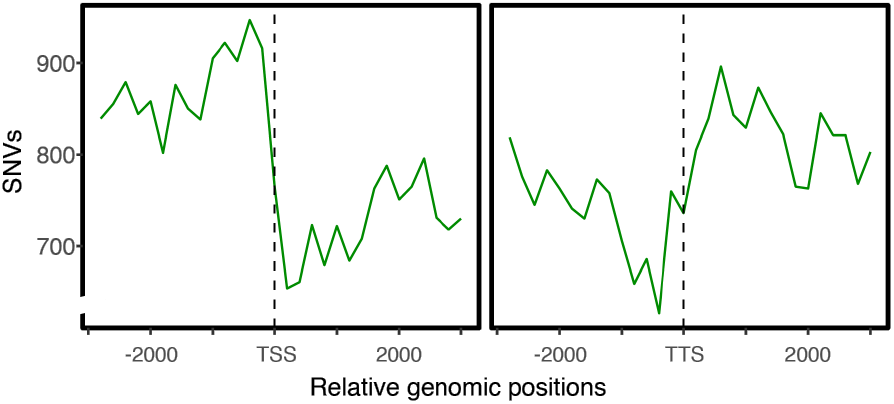
Confirmation of reduced mutation rate in gene bodies with a rice fast-neutron mutation data set. Distribution of *de novo* mutations in 1,504 fast-neutron-treated rice lines, with >99% validation rate of SNVs [43]; for further details see [44].

### Population genetic theory confirms the evolutionary potential of targeted DNA repair

Liu and Zhang [13] also posit that the mutation biases observed in [1] violate the drift barrier, which limits the evolution of intragenomic mutation rate heterogeneity [3]. We propose that this is not the case and that any apparent conflict with the drift-barrier hypothesis can be reconciled under the emerging consensus on mechanisms of targeted DNA repair, increasingly recognized to be enabled by binding of repair proteins to histone modifications linked to gene regulation, such as MSH6 binding H3K36me3 in transcribed exons via PWWP in vertebrates [6,8,9,12]. Chromatin features are not independent of gene function and they can simultaneously serve as targets for elevated DNA repair. The evolution of chromatin-linked targeted DNA repair can therefore overcome the barrier imposed by drift, because it affects the mutation rates of thousands of loci, which in turn constitute a suitably large target for selection. These mechanisms can promote DNA repair specifically in functionally constrained regions of the genome, increasing the specificity of repair in regions where mutations are more likely to be deleterious. This was first shown by Martincorena and Luscombe [4] and expanded on in [1]. In *A. thaliana,* H3K4me1 is the most highly enriched feature of coding sequences, especially in constrained genes (Fig. 4). In humans, H3K36me3 is the most highly enriched feature of coding sequences, and is especially enriched in genes subject to the strongest constraints [4,45].

**Figure 4.**
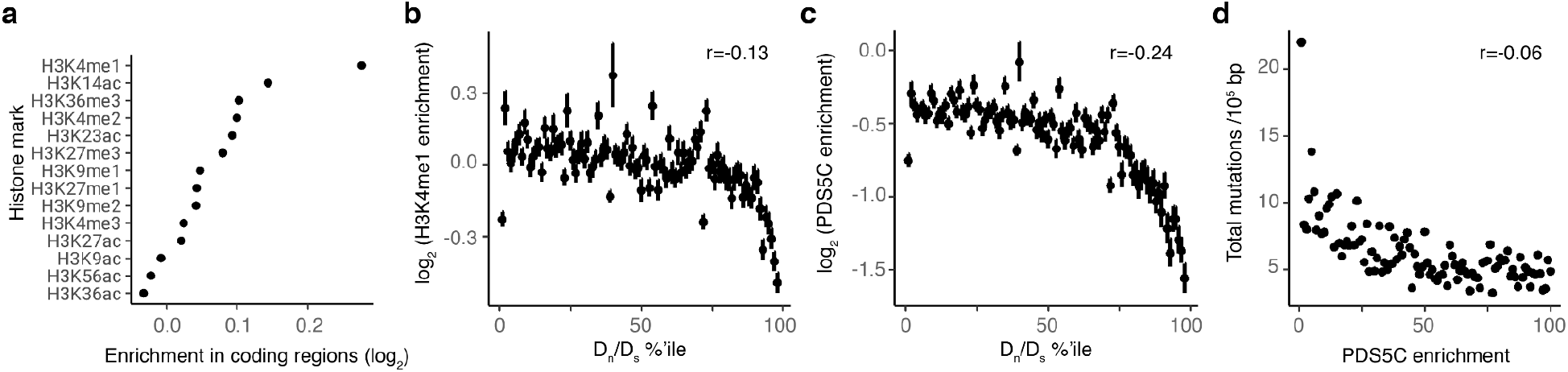
Enrichment of H3K4me1 and DNA repair in evolutionarily constrained regions. **(a)** H3K4me1 is the most highly enriched histone modification found in coding sequences relative to other genome regions in *A. thaliana.* Data from the Plant Chromatin State Database [46]. Enrichment on the x-axis corresponds to the ratio of the observed to the expected presence under uniformity. **(b)** H3K4me1 is enriched in coding regions of genes subject to stronger constraint, estimated by D_n_/D_s_. **(c)** PDS5C, a repair protein that binds H3K4me1 via its Tudor domain, is enriched in coding regions of more constrained genes. PDS5C ChIP-seq data were from [40]. Enrichment was calculated as log_2_[(1 + n_ChIP)/N_ChIP] – log2[(1 + n_Input)/N_Input)], where n_ChIP and n_Input represent the total depth of mapped ChIP-seq and input fragments in a region, respectively, and N_ChIP and N_Input represent the total depth of mapped unique fragments. **(d)** Germline mutations, aggregated from several independent datasets [1,15–17,19–21,41,42], are more frequent in genes depleted for bound PDS5C. r in **b**, **c**, **d** corresponds to Pearson’s Correlation Coefficient, with P < 2e-16 for all three. Error bars indicate standard error of the mean.

The drift barrier exists when N_e_ * u * s * d_u_ * p_d_ * L_segment_ < 1, where *N_e_* is the effective population size, *u* is the mutation rate, *s* is the average selection coefficient on deleterious mutations, *d_u_* is the degree of change in mutation rate, *p_d_* is the proportion of sites subject to purifying selection, and *L_segment_* is the absolute size of the affected genomic regions [3]. In the case of chromatin targeted DNA repair of constrained genomic regions, both *L_segment_* and *p_d_* are elevated. Liu and Zhang [13] suggest that this cannot be satisfied for the evolution of gene-specific modifiers in species with moderate population sizes. However, we posit that this argument does not apply to the mechanisms and mutation biases in question, which reflect genome-wide systems affecting at the same time genomic regions that are many orders of magnitude larger than single genes. In [1], it was shown that the drift-barrier can be overcome as soon as only 1.5% of the portion of the *A. thaliana* genome is preferentially targeted for DNA repair by a specific chromatin mark such as H3K4me1. Not only does the drift-barrier therefore readily accommodate chromatin-mediated repair mechanisms, the underlying model also provides quantitative population genetic evidence that explains how known mechanisms of targeted DNA repair can and have evolved.

## DISCUSSION

Here, we have revisited analyses and conclusions of the study [1] that has come under scrutiny by a recent publication from Liu and Zhang [13]. The primary results reported in [1] remain similar after more stringent filtering of variant calls (Fig. 1), with analyses of germline mutations alone, aggregated from several independent sources, supporting several core conclusions that were previously based to a large extent on somatic mutation calls. An approximately 50% reduction of mutation rates was observed in gene bodies in [1], which is confirmed here with germline mutations only from multiple independent mutation accumulation experiments in *A. thaliana* (Fig. 2a, b) [15–17,19–21,41,42]. An additional reduction of ~30% was observed in essential genes. This is also confirmed here with the new analyses of germline mutations only (Fig. 2c), and it holds even when analyses are restricted to non-coding genic sequences.

In addition, new results point to potential underlying biochemical mechanisms that can explain mutation bias in *A. thaliana.* Proteins with Tudor domains, as they are found in some plant DNA repair proteins, specifically bind to regions with lower mutation rates (Fig. 4d) [40,44]. A potential mechanistic basis (Fig. 5) comes from ChIP-seq data, which indicate that plant Tudor domains preferentially bind regions marked with H3K4me1 [40]. Both plant PDS5, central to homology-directed repair, and plant MSH6, central to mismatch repair, contain such Tudor domains [44]. This is reminiscent of human repair proteins, including MSH6, which are targeted to gene bodies via PWWP domains that bind H3K36me3 [6,8,9,12] (Fig. 6). Evidence of DNA repair proteins binding to regions with lower mutation rates cannot be reasonably explained by sequencing artifacts. That targeted DNA repair, as opposed to better protection of DNA from damage, is likely responsible for reduced mutation rates in genic sequences is supported by the finding that mutation rates are specifically increased in gene bodies of *msh2* knockout plants (Fig. 2b). MSH2 functions as a dimer with MSH6, which is likely targeted to H3K4me1 via its Tudor domain [10,44].

**Figure 5.**
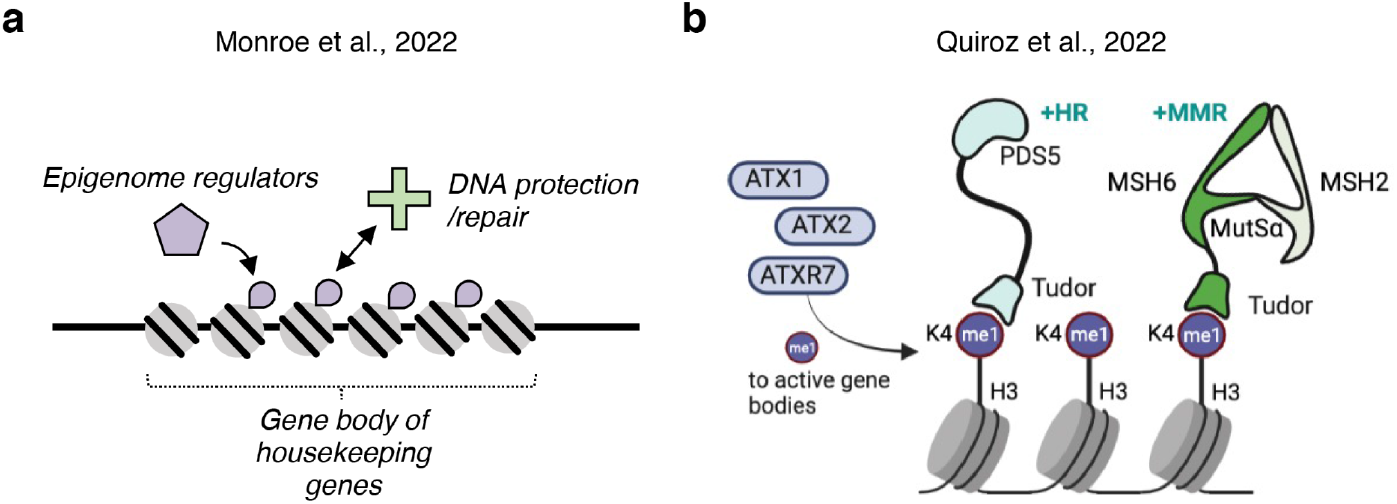
Mechanistic models of mutation bias reflecting natural selection. **(a)** General mechanism proposed in [1]. **(b)** Updated knowledge of biochemical mechanisms underlying reduced mutation rates in gene bodies established by recent discoveries in plants and synthesized in [44]. HR, homology directed repair; MMR, mismatch repair.

Additional recent studies that support the core findings of [1] include reports of (i) reduced germline and somatic mutations rates in transcribed genes, in genome regions marked by epigenomic features characteristic for gene activation, and in gene bodies, observed in several algae, plants as well as humans and nematodes [17,21,36,42,47–52]; (ii) relationships between histone modifications – especially those linked to gene activation –, mutation rates and DNA repair across diverse species [51,53–60]; and (iii) H3K4me1 affecting rates of CRISPR-mediated mutagenesis in plants [61]. That the role of histone modifications in driving DNA repair localization is well-established is also reflected in cancer therapeutics targeting histone states being widely pursued (e.g., [62–66]).

Another important question is how universal mutation bias is. Transcription is mutagenic [68], and the greater DNA damage caused by higher average transcription levels of constrained genes may lead to higher mutation rates in the human germline [69]. How well the effects of increased mutation due to transcription are compensated by transcription-coupled DNA repair may depend on transcription rate [70–72].

Of note, Boukas and colleagues [45] recently leveraged a very large *de novo* mutation data set [73], finding that more evolutionary constrained genes have a lower rate of synonymous mutation in humans, which is consistent with at least one previous report [74]. Specifically, these authors reported that the synonymous mutation rate of genes under least constraint, as determined from their LOEUF (“loss-of-function observed/expected upper bound fraction” [75]) decile, is almost twice as high as that of genes under the strongest constraint [45]. In contrast, Liu and Zhang [13] report that the transcribed regions of genes under greater evolutionary constraint have higher total mutation rates in humans than genes under lower constraint. It is possible that this discrepancy is a reflection of not accounting for constrained genes containing more intronic sequence, which could be more prone to mutation [12,36,76]. Alternatively, Liu and Zhang [13] may not have accounted for zero-inflated data, which can affect mutation rate estimates [72].

Boukas and colleagues showed that epigenomic marks can be under selection, and they propose that the lower mutation rate of evolutionary constrained genes is driven to a large extent by H3K36me3, a major target of DNA repair in humans via proteins with PWWP domains [45]. This parallels findings in *A. thaliana* and rice, where H3K4me1 marks constrained genes and at the same time is important for recruiting DNA repair proteins [1,44,61]. Liu and Zhang [13] also do not find mutation bias in yeast. Yeast MSH6 lacks a histone reader domain, which distinguishes the yeast protein from its plant and vertebrate homologs (Fig. 6), which may explain the absence of DNA repair being targeted to evolutionary constrained sequences by epigenomic features in this taxon. Clearly, more studies are needed to understand the diversity in mutation biases and their underlying mechanisms.

**Figure 6.**
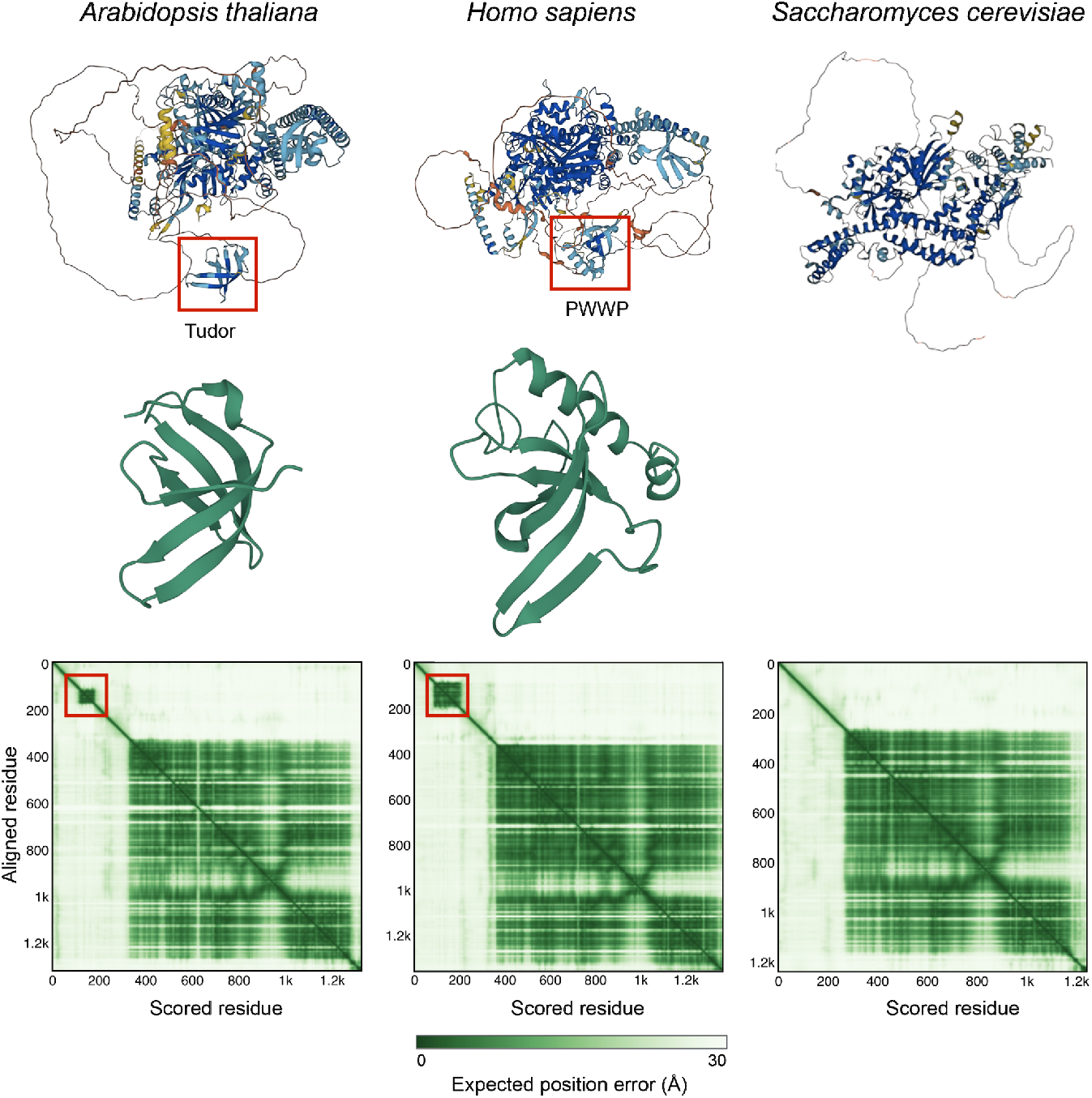
Structure of DNA mismatch repair protein MSH6 from different species. Top, AlphaFold structures [67], with Tudor and PWWP domains in *A. thaliana* and *H. sapiens* highlighted in red, and close-ups of Tudor and PWWP domains below. Bottom, confidence of structural prediction for repair domains, which make up the bulk of the proteins, as well as Tudor and PWWP domains, which are highlighted in red. An analogous, structured chromatin binding domain outside the conserved repair domain is not apparent in *S. cerevisiae.*

In summary, the main conclusions of [1] remain supported after removal of likely homopolymer errors or excluding C:G>T:A mutations, which are particularly frequent in TEs. Additional lines of evidence for the conclusions of [1] include a meta-analysis of germline mutations from independent sources. We nevertheless note that most of the analyses in [1] used both germline and somatic mutation calls, weighing both types of evidence equally. A more conservative approach would have been to account for the substantially greater uncertainty associated with somatic mutation calls. The models of mutation probability, which are at the core of [1], are least affected by such considerations, since they relied to a large extent on germline calls. However, the testing of epigenome-driven mutation probability predictions with empirically observed mutations, including the ones shown in Fig. 2b, 3c, and 4c of [1], were overwhelmingly based on putative somatic mutation calls. The much greater uncertainty associated with these calls should have been acknowledged. This concern is mitigated by new analyses that only use germline mutations (Fig. 2, 3, and 4d of the current work) and which support the inferences from [1].

We therefore believe the hypothesis that deleterious mutation rate can be reduced by links between the epigenome and mutation rate via targeted DNA repair [1] to be correct because it is supported by independent datasets, confirmed with more high-quality datasets than those used in [1], because evolutionary and population genetic analyses in other systems point in the same direction [45], and because a potential mechanistic basis is becoming better understood, as discussed throughout this study. At the same time, it is clear that the original evidence in [1] for the hypothesis of reduced mutation rate in constrained regions of the genome was not as strong as we thought when submitting the work for publication, because the much higher error rate in somatic mutation calling was not fully accounted for. The conclusions of [1] would have been strengthened by the new analyses presented here.

## METHODS

### Recalling and filtering somatic mutation data in training set for mutation probability model in [1]

For the mutation probability model in [1], variant calls from 107 MA lines [16] were used. Here, the Illumina sequencing reads (NCBI accession SRP133100) were processed using fastp [77] to remove low quality reads (<Q15) and to trim adapters. Clean reads were aligned to the *A. thaliana* telomere-to-telomere reference genome [29] using BWA mem (0.7.17-r1188) [78] and NextGenMap (0.5.5) [79] with default parameters. Duplicate reads were marked GATK (MarkDuplicates, version 4.2.3.0) [80]. Variants for each accession were called with GATK (HaplotypeCaller). All gVCF files were merged using GATK (CombineGVCFs) and genotyped using GATK (GenotypeGVCFs).

After mapping and variant calling, variants were removed if reads supporting the alternative allele were observed in more than 1 sample, mapping quality was <60, variants reflected potential homopolymer errors (defined as variants in which an alternative allele is adjacent to a repeat of at least 3 bases of the same type [G, A, T, C] [22]), the distance to a neighboring mutation is <10bp, or if the alternate allele was only supported by reads from one strand.

### Mutations from independent datasets

Mutations were obtained from 10 mutation accumulation datasets in *A. thaliana,* originally reported in [1,15–21,41,42]. If not reported in the original study, gene body mutations were counted by whether a mutation occurred in the regions of the TAIR10 genome annotated as protein coding genes. The fraction of the genome corresponding to gene bodies (transcribed regions) was 50.4%.

### Gene-level estimates of constraint, repair, chromatin state, and mutation rates

Chromatin marks, essentiality, lethality, frequency of loss-of-function alleles in natural populations from the 1001 Genomes (1001G) Project, D_n_/D_s_, P_n_/P_s_, number of tissues expressed, sequence diversity in 1001G natural populations, and variation in expression have been reported in [1], derived from original sources [30,46,81–85].

We compared germline mutation rates aggregated across independent studies between the upper and lower 50% of genes according to quantitative measures of constraint. The total number of mutations and total sequence space was compared by χ-squared tests. A 95 % confidence interval was generated by bootstrapping. In our reanalysis of germline mutations across studies we also re-ran tests restricted to non-coding genic sequences only, to ensure that the conclusions were not driven by cryptic selection on coding sequences in the mutation accumulation experiments. We also re-ran analyses after removing regions with zero mutations, to control for potential biases introduced by including zero-inflated data [72]. Relevant code is maintained at https://github.com/greymonroe/mutation_bias_analysis2 and https://github.com/greymonroe/polymorphology.

PDS5C ChIP-seq data have been published [40] (GEO Accession GSE154302) and were analyzed with the script https://github.com/greymonroe/mutation_bias_analysis2/blob/main/code/PDS5C%20chip.R.

## AUTHOR CONTRIBUTIONS

All authors contributed to this study. Kevin D. Murray and Wenfei Xian were not among the authors of [1], the study that has come under scrutiny by the analyses in [13].

## ACKNOWLEDGMENTS

We thank members of the Weigel lab for discussion. This work was supported by a Marie Skłodowska-Curie Postdoctoral Fellowship (K.D.M.), by startup funds from the University of California Davis (J.G.M), by NSF grants DEB 0844820 and DEB 1257902 (C.B.F.), and by DFG grant ERA-CAPS 1001G+ and the Max Planck Society (D.W.). Research was conducted at the University of California Davis, which is located on land which was the home of the Patwin people for thousands of years.

## SUPPLEMENT

In [1], we had reanalyzed data from [31]. We inadvertently mapped the same reads twice to the genome (bwa mem R1 R1). A consequence was that singletons, that is, variants supported by only on read, were called as non-singleton variants. These are expected to include an undue number of errors, which likely exist as singletons, along with low-frequency somatic mutations, such as those that arise late in leaf development. Unfortunately, we cannot distinguish these two categories, which greatly reduces confidence in these data.

We recalled variants after correcting the mapping pipeline to match the methods described in [1] (Extended Data Fig. 4c in [1]). We observe the same overall patterns: lower mutation rates in gene bodies (Fig. 2b in [1]), differences in mutation rates according to annotation of genes (Fig. 3c in [1]), correlation between quantiles of observed mutation rates per gene and population measures of evolutionary constraint (Fi. 4c in [1]), and a relationship between epigenomic and other features on the one hand and mutation rates of lethal-effect and constitutively expressed genes on the other hand (Extended Data Fig. 8b,d in [1]).

We note that comparison of the original calls of this specific dataset in [1], which included singletons, with the corrected data set indicates that calls of singletons include fewer homopolymer bleed-over artifacts. In the original analysis only 10% of single-nucleotide variants were possible homopolymer bleed-over errors (ALT variant same as neighboring homopolymer), whereas in the re-called variants 34% of single-nucleotide variants are potential homopolymer bleed-over errors. Thus, singletons may contain a lower fraction of artifactual calls. Therefore, while these corrected data may more closely reflect the methods described in [1], the resulting variants are not necessarily more accurate in terms of their distribution in relation to true mutation rates across the genome, based on the expectation that the overwhelming majority of *de novo* somatic mutations is found in single or very few cells, given that meristematic cells – in which somatic mutations would result in many cells with shared mutations – have elevated DNA repair compared to differentiated cells [86,87].

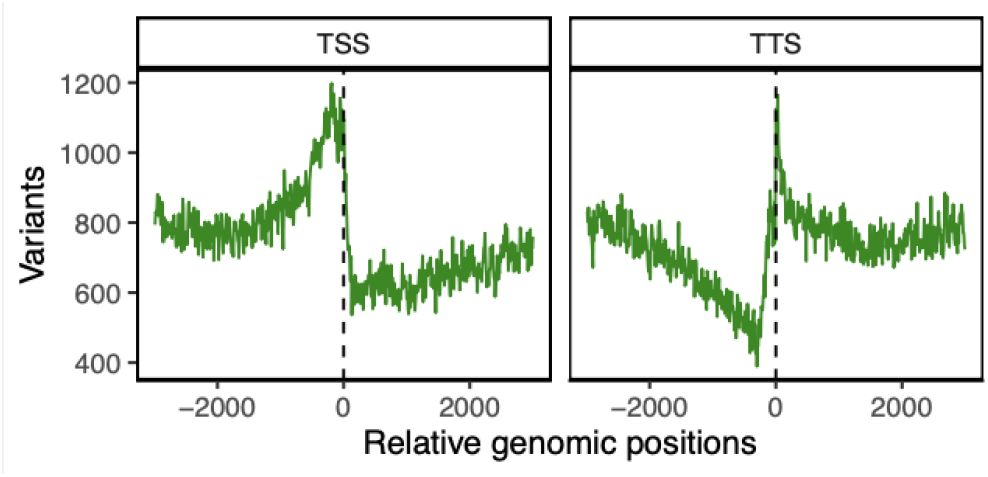
**Corrected Figure 2b of** [1]. Observed de novo mutations from all independent mutation accumulation datasets (mean ± 2 s.e.m. in grey, bootstrapped). TSS, transcription start site; TTS, transcription termination site. Coordinates in bp.

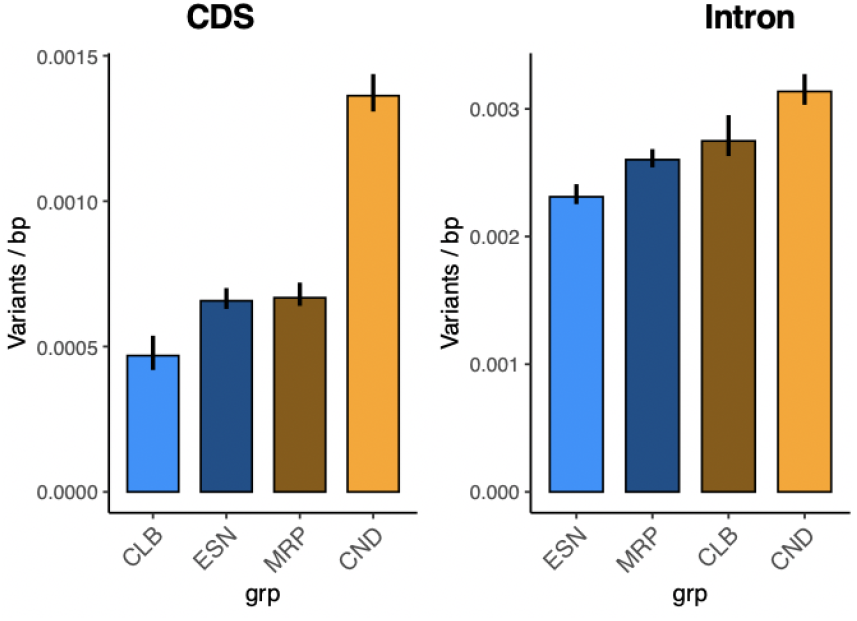
**Corrected Figure 3c of** [1]. Total observed mutation rate (±2 s.e.m., bootstrapped) in genes (n = 2,339) with experimentally determined functions. The bars are coloured according to relative differences in mutation rates among genes classified by function (that is, orange refers to high mutation rate and blue represents low mutation rate). ESN, essential genes; CLB, cellular and biochemical phenotypes; CND, environmentally conditional phenotypes; MRP, morphological phenotypes.

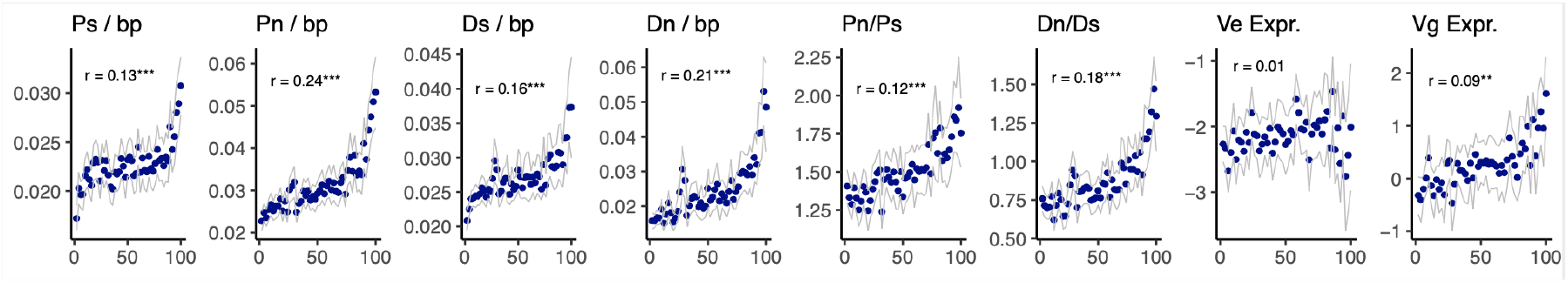
**Corrected Figure 4c of** [1]. Rate of sequence evolution across quantiles of observed mutation rates. Synonymous (Ps) and non-synonymous polymorphism (Pn) in natural populations, synonymous (Ds) and non-synonymous divergence (Dn) from *A. lyrata,* environmental variance of gene expression (Ve expr.), genetic variance of gene expression (Vg expr.).

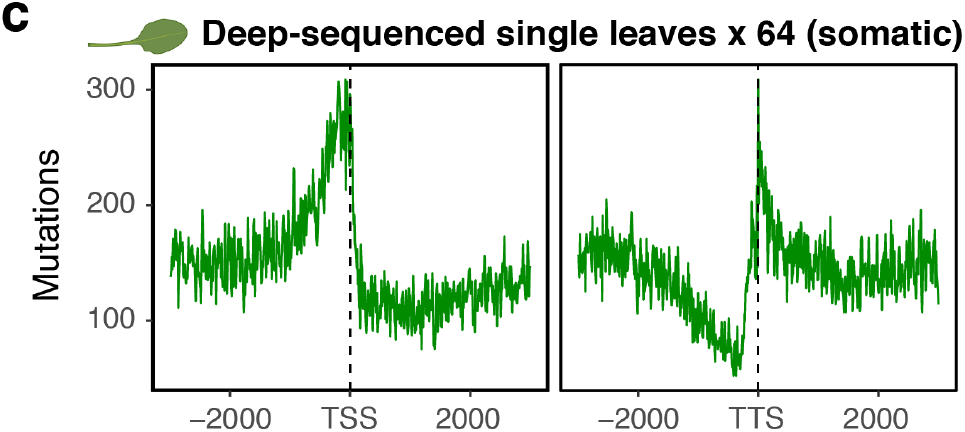
**Corrected Extended Data Figure 4c of** [1]. Somatic variants detected from reanalysis of 64 individual leaves from two Col-0 plants [31].

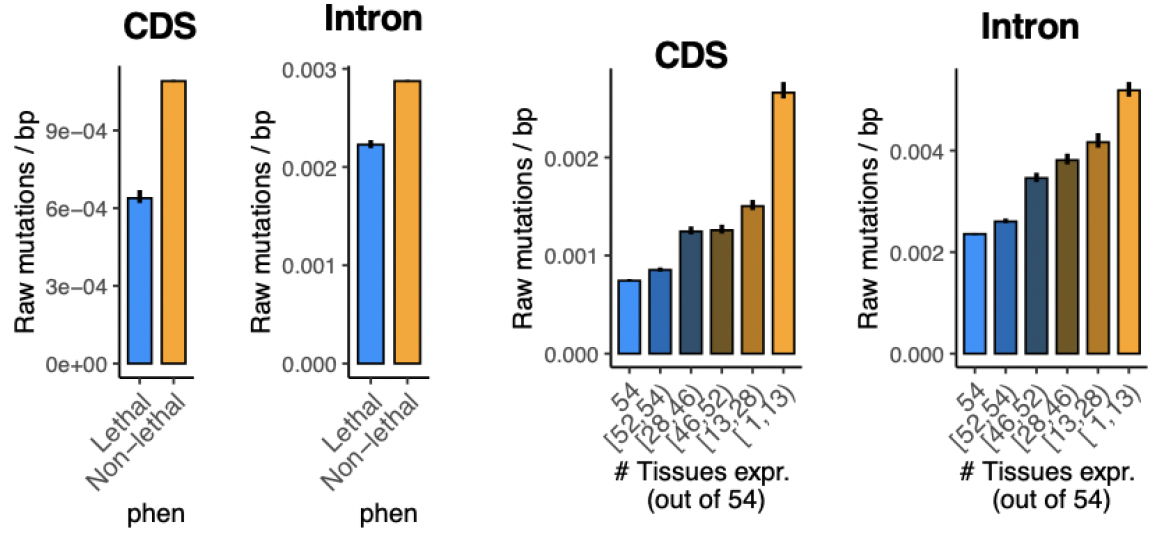
**Corrected Extended Data Figure 8b, d of** [1]. **b**, Total mutation rate (±2 s.e.m., bootstrapped) in lethal- and non-lethal-effect genes (n = 27,206 genes). **d,** Total mutation rate (±2 s.e.m., bootstrapped) in genes binned according to the number of tissues in which they are expressed (n = 25,987 genes with tissue-specific expression data).

